# Characterization of exosporium layer variability of *Clostridioides difficile* spores in the epidemically relevant strain R20291

**DOI:** 10.1101/2020.04.05.024893

**Authors:** Marjorie Pizarro-Guajardo, Paulina Calderón, Alba Romero-Rodriguez, Daniel Paredes-Sabja

## Abstract

*Clostridioides difficile* is a Gram-positive anaerobic intestinal pathogenic bacterium and the causative agent of antibiotic-associated diarrhea and spores are the transmission vehicle of the disease. In *C. difficile* spores, the outermost exosporium layer is the first barrier of interaction with the host and should carry spore ligands involved in spore-host interactions. *C. difficile* forms two types of spores (i.e., thin and thick exosporium layers). In this communication, we contribute to understand several biological aspects of these two exosporium morphotypes. By transmission electron microscopy, we demonstrate that both exosporium morphotypes appear simultaneously during sporulation and that the laminations of the spore-coat are formed under anaerobic conditions. Nycodenz density-gradient allows enrichment of spores with a thick-exosporium layer morphotype and presence of polar appendage. Using translational fluorescent fusions with exosporium proteins BclA3, CdeA, CdeC and CdeM as well as with several spore coat proteins, we observed that expression intensity and distribution of SNAP-translational fusions in R20291 strain is highly heterogeneous. Electron micrographs demonstrate that multicopy expression of CdeC, but not CdeM, SNAP translational fusion, increases the abundance of the thick exosporium morphotype. Collectively, these results raise further questions on how these distinctive exosporium morphotypes are made during spore formation.

## Introduction

*Clostridioides difficile* is the leading cause of nosocomial antibiotic-associated diarrhea [1, 2]. Recurrence of CDI is a major problem with an incidence of 25 to 65% of patients, associated to increase severity of symptoms and time in hospital settings [3]. The main factor involved in the recurrence of CDI is the formation of metabolically dormant spores during the infection [4]. These newly formed spores are essential for the persistence of *C. difficile* in the host and the transmission of the disease to a new susceptible host [4]. Sporulation is initiated by an asymmetric cell division that leads to the formation of the mother cell and the incipient forespore [5]. Later, the mother cell engulfs forespore and mediates the assembly of structural layers, including spore peptidoglycan cortex, inner and outer coat, and exosporium. Finally, the mother cell lyses and releases the mature spore [5]. Sporulation in *C. difficile* is asynchronous, therefore at a given time, multiple stages of development can be found [6], thus hampering the study of synchronized cultures.

The outermost surface of *C. difficile* spores is thought to play an important role in host-spore interaction by ligands involved in interactions between spore and host cellular receptors [7-9]. Recent evidence indicates that *C. difficile* produces spores with two distinctive morphotypes: that is, an exosporium layer with a thin electron dense layer, and an exosporium layer with a thick electron dense layer forming prominent bumps [10, 11]. By sonication and trypsin digestion of the spore surface, enriched fractions of exosporium peptides revealed the presence of several proteins which were shown to be unique for this outer most layer [12]. These include, three cysteine-rich proteins (CdeA, CdeC and CdeM) and three collagen-like proteins (BclA1, BclA2 and BclA3) [12, 13]. Some spore coat proteins, including CotA, CotB, CotD and CotE were also observed, most likely as part of the spore-coat surface that interacts with the exosporium layer [12]. The spore coat and exosporium layer are formed during late stages of sporulation, under the control of sigma factors SigE and SigK [5], which regulate the expression of the aforementioned proteins. However, the mechanisms that govern exosporium assembly remain unclear.

To date, two morphogenetic cysteine-rich proteins, CdeC and CdeM, have been identified as essential for the correct assembly of the exosporium layer [8, 14, 15]. Both are required for the location of several spore-coat and exosporium proteins to the outer spore surface layers [8] and the absence of CdeM seems to be involved in the conformation of electron dense material in the spore surface [14]. How these proteins contribute to the variability observed in the outermost layer of *C. difficile* spores remains unclear. In this work, we explore new features in exosporium layer variability. We used transmission electron microscopy to demonstrate that thick and thin exosporium spores are produced simultaneously during sporulation of *C. difficile*. By a density gradient, spores can be separated and an enriched fraction in thick-exosporium spores and appendage containing spores can be obtained. By the use of a fluorescence reporter, we observed high heterogeneity in fluorescence intensity and distribution of protein-SNAP fusions. Notably, CdeC-, but not CdeM-SNAP translational fusion, leads to an increase in the relative abundance of thick exosporium spores and exacerbated thick exosporium layer. Collectively, these results provide more evidence of the high variability of *C. difficile* spore outermost layer.

## Materials and Methods

### Spore purification of *C. difficile* R20291

*C. difficile* strain R20291 used in this study was routinely grown under anaerobic conditions on BHIS broth (3.7% Brain Heart Infusion supplemented with 0.5% yeast extract, 1% cysteine). 16 h cultures were diluted to 1:500 dilution and 100 μl were plated onto 70:30 (63 g Bacto peptone, 3.5 g protease peptone, 11.1 g BHI medium, 1.5 g yeast extract, 1.06 g Tris, 0.7 g NH4SO4, 15 g agar per litter) or TY (3% Trypticase Soy-0.5% yeast extract) agar plate and incubated at 37°C for 7 days. After incubation, plates were scraped up with 1 ml of ice-cold sterile water. Spores were gently washed five times (resuspension in ice-cold sterile water and centrifugation at 16,000 × g for 5 min). Next, spores were gently loaded onto a 1.5 ml tube containing 300 μl of 45% Nycodenz solution and centrifuged for 40 min at 16,000 × g. After centrifugation, the supernatant was removed, and the spore pellet was washed five times. The spores were counted in a Neubauer chamber and concentration adjusted at 5×10^9^ spores/ml to store at −80°C until use.

### Transmission electronic microscopy (TEM)

Spores (2×10^8^) were fixed with 3% glutaraldehyde on a 0.1M cacodylate buffer (pH 7.2), incubated overnight at 4°C, and stained for 30 min with 1% tannic acid. Samples were further embedded in a Spurr resin [16]. Thin sections of 90 nm were obtained with a microtome, placed on glow discharge carbon-coated grids and double lead stained with 2% uranyl acetate and lead citrate. Spores were analyzed with a Philips Tecnai 12 Bio Twin microscope at Unidad de Microscopía Avanzada in Pontificia Universidad Católica de Chile.

### Separation by Nycodenz-gradient of *C. difficile* R20291 spores

A Nycodenz-gradient was performed in which 64% Nycodenz was layered in the bottom followed by layers decreased in 1% until 56%. Spores (2.5×10^9^ spores) were gently added to the surface and centrifuged at 5,000 × g for 40 min in a swim bucket rotor. Two fractions were formed, which were recovered in aliquots of 1 ml by removing them gently from the surface of the Nycodenz gradient. Collected fractions were washed with 1 ml of sterile water and adjusted to a final concentration of 5×10^9^ spores/ml.

### Adherence of fractionated *C. difficile* R20291 spores

Caco-2 cells (ATCC, U.S.A) were used for infection with separated spores at 2- and 8-days post confluence. Spores were added at a MOI of 10 and incubated for 3 h at 37°C under aerobic conditions with 5% CO2 [7]. Unbound spores were rinsed off with three washes of PBS. Unwashed wells were employed for total spore count. Caco-2 cells lysed in 80 μl 0.06% Triton X-100 for 30 min at 37°C. Cell-spore lysate were serially diluted, plated on BHIS-0.1% Sodium Taurocholate, and incubated in anaerobic conditions at 37°C for 48 h. The number of colony forming units (CFU) per ml was determined, and the percentage of adherence was calculated using the relation: (CFU ml^-1^/TOTAL CFU ml^-1^) × 100. The data represents the averages of the results of three independent experiments.

### Protein-SNAP fusion construction

According to the reference annotated genome of *C. difficile* strain R20291 (FN545816), genes and their coding regions used in this work were *cdeC* (CDR20291_0926), *cdeM* (CDR20291_1478), *bclA3* (CDR20291_3193), *cotA* (CDR20291_1511), *cdeB* (CDR20291_2642), *cotB* (CDR20291_1360), *cotE* (CDR20291_1282), *cotD* (CDR20291_0523) and their respective promotor region were independently cloned in plasmid pFT58. Briefly, promotor and coding region sequence were amplified with primers listed in Table S1, and cloned between EcoRI/BamHI sites in pFT58, which contains sequence for SNAP. Plasmid were stored in DH5*α*. Constructed plasmids are listed in Table S2.

### Fluorescent microscopy and analysis of SNAP fusion

Constructed plasmids (Table S2) were conjugated into *C. difficile* R20291 using *E. coli* CA434 as the donor strain. Transconjugant *C. difficile* colonies were selected for resistance to 15 mg/ml thiamphenicol. *C. difficile* R20291 strains carrying SNAP fusion vectors were grown in BHIS - thiamphenicol, seeded in 70:30 at 1:100 dilution and incubated for 48 h at 37°C. A fraction of the plate was scrapped and resuspended in PBS-250 nM of SNAP-Cell® Oregon Green® (Invitrogen) and incubated for 30 min at 37°C [17]. Finally, labeled sample were pelleted and resuspended in PBS containing 10 ng/ul FM4-64 and 2.4 μM Hoechst, incubated at room temperature for 2 min and centrifugated 5 min at 6000 × g. The pellet was resuspended in 20 ul of PBS and 5 μl of the sample was mounted in coverslips with agarose pad and observed with BX53 Olympus fluorescence microscope. The fluorescence images were analyzed with ImageJ. At last, 200 spores were analyzed in three independent experiments.

## Results

### Thin and thick exosporium spores are observed simultaneously in sporulating cultures of *C. difficile*

We sought to determine whether the thick and thin exosporium morphotypes were formed simultaneously during spore formation. Sporulating cultures of 70:30 agar plates, were fixed under anaerobic conditions to assess whether the laminations of the spore-coat and exosporium thickness occurs during spore-development and not after aerobic processing of the sporulating culture. Transmission electron micrographs from anaerobically fixed cultures show at early sporulation stage (1-day-old sporulation culture) several mother cells containing mature spores with both exosporium morphotypes (Fig. 1A). We also observed that coat laminations can be clearly observed as soon as 24 h of sporulation (Fig. 1B). At 1 day-old 70:30 sporulating cultures, percentages of thin and thick exosporium layer morphotypes are 84 and 16%, respectively (Fig. 1B). In 5-days-old 70:30 sporulating cultures, the percentage of thin and thick exosporium morphotype spores was 76 and 24%, respectively (Fig. 1B). Similar results were observed in TY agar media (Fig. S1). In TY medium, 1-day-old sporulating cultures revealed the presence of vegetative cells at different stages of the developmental process (i.e., asymmetrically divided cells, engulfed forespore and mature endospore) (Fig. S1A). Collectively, these results support the notion that during *C. difficile* sporulation spores with a thick or thin exosporium layer are formed simultaneously during sporulation.

**Figure 1.**
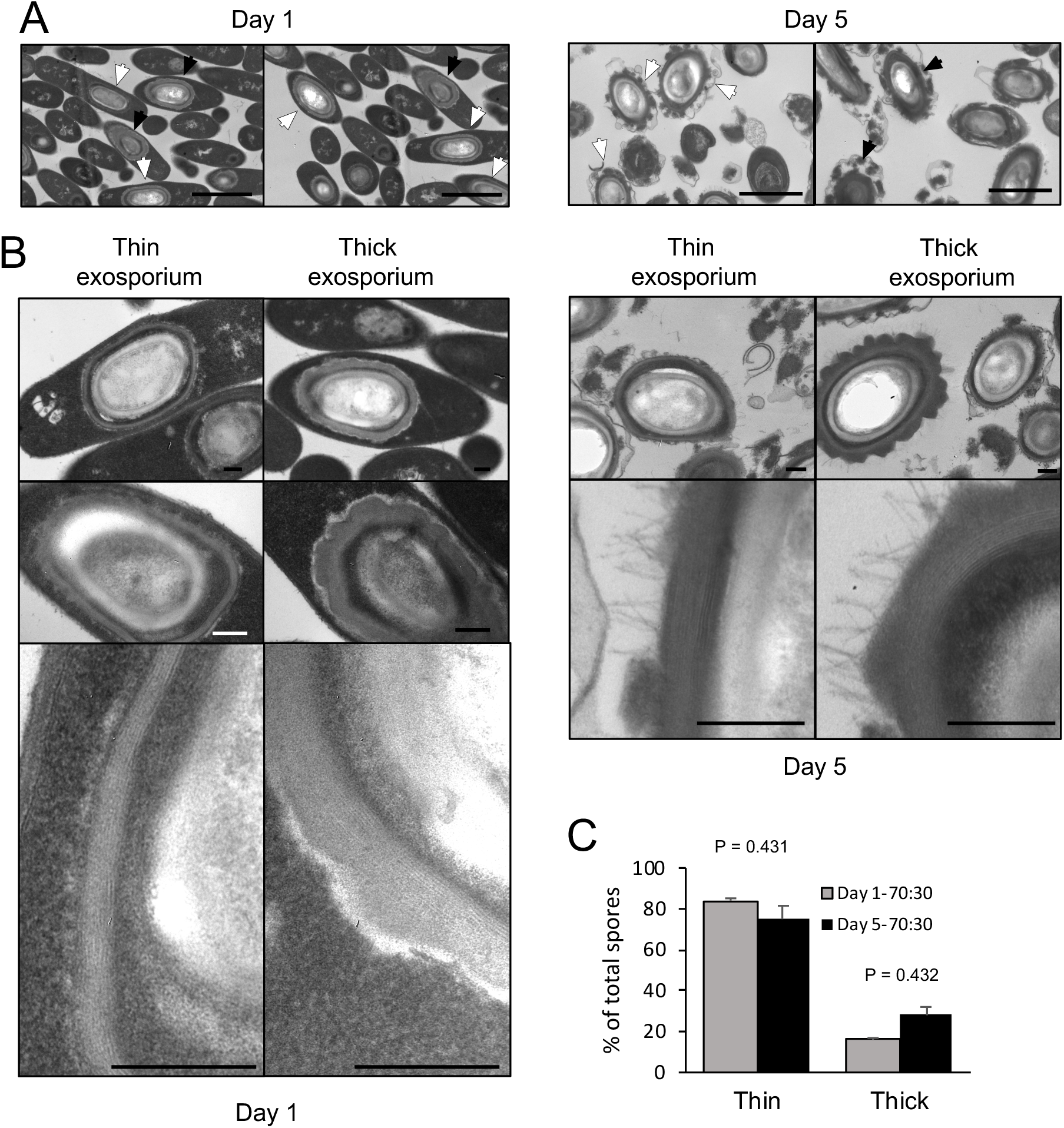
Transmission electron micrographs of 1-day- and 5-day-old sporulating cultures in 70:30 agar plates. A) Sporulating cultures in 70:30 medium harvested after 1 day (left) and 5 days (right) of growth in 70:30 medium. Cultures were fixed under anaerobic conditions prior to processing for transmission electron micrograph analysis. White arrow: thin exosporium spore, Black arrow: thick exosporium spore. Bar scale: 2 μm. B) Ultrastructural analysis of the exosporium layer. Thin and thick exosporium morphotype spores evidenced at day 1 (left panel) and day 5 (right panel). Scale bar: 200 nm. C) Percentage of thin and thick exosporium at day 1 (gray bar) and day 5 culture (black bar). Bar indicates standard error obtained with 3 different batch of culture fixed, processed and analyzed separately. N = 50 spores. P-values indicated.

### Enrichment of polar appendage and thick exosporium morphotypes by Nycodenz gradient of *C. difficile* spores

The simultaneous appearance of both exosporium morphotypes during sporulation in different sporulating media (70:30 and TY) raised the question of whether *C. difficile* spores can be separated into two populations according to their exosporium thickness. Herein, we used a Nycodenz gradient to fractionate TY pure spores and observed 2 fractions of spores were the upper fraction stacked at 58% of Nycodenz and the lower fraction stacked at 62% of Nycodenz (Fig. 2A). Transmission electron analysis revealed that while the upper fraction maintains the same proportion of thick (58% of spores) and thin (42% of spores) exosporium spores as unfractionated TY spores (Fig. S1), more than 80% of the spores analyzed of the lower fraction were thick exosporium spores (Fig. 2B).

**Figure 2.**
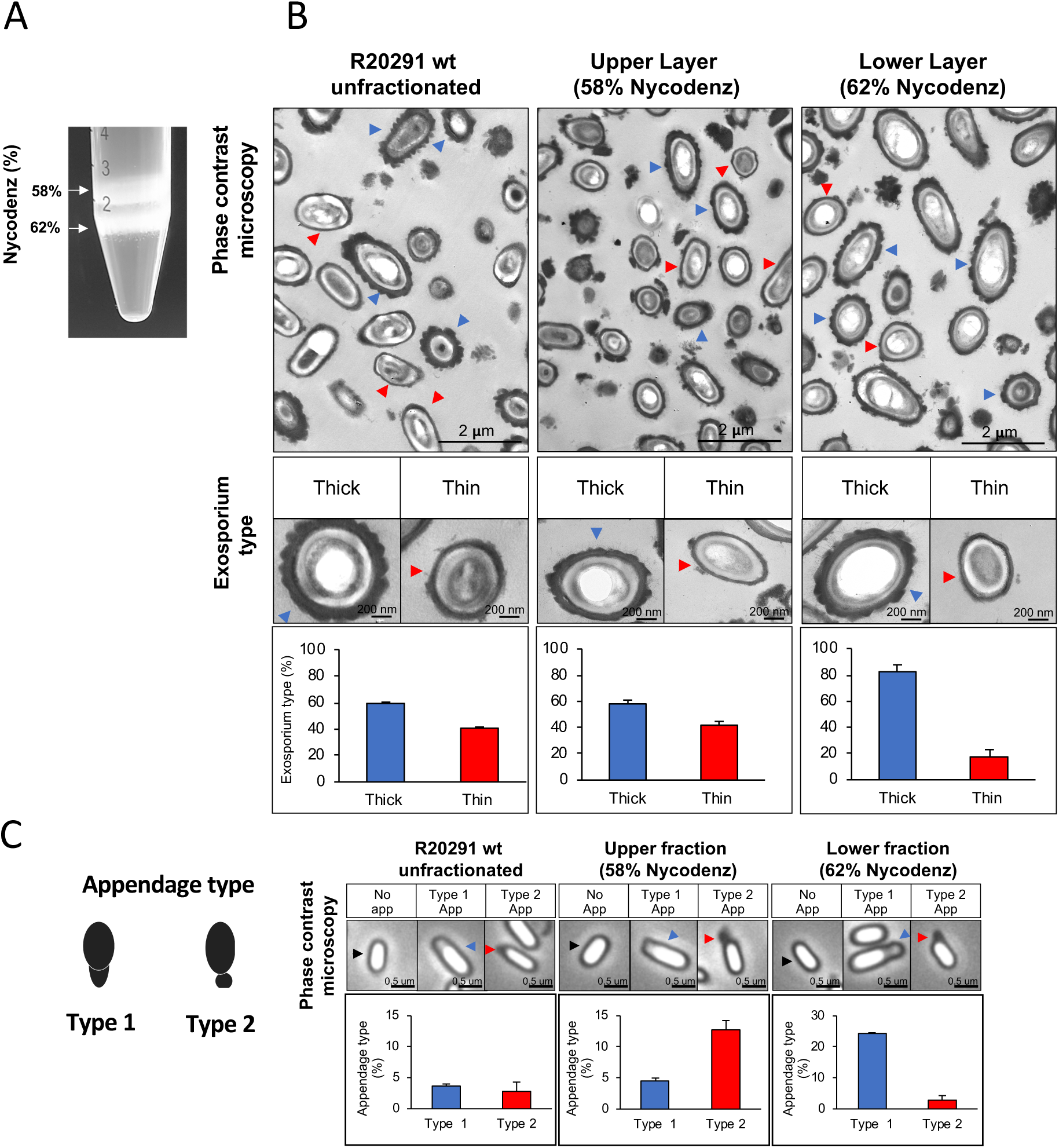
*C. difficile* R20291 spores separated in Nycodenz gradient. A) Spores were separated with a Nycodenz gradient from 56% to 65% by centrifugation at 5,000 × g for 40 min. Two-fractions were obtained: fraction (58% Nycodenz) and lower fraction (62% Nycodenz). White arrows indicate both fractions. B) Classification according to exosporium thickness of unfractionated, upper fraction, and lower fraction spores. Blue arrowheads indicate spores with a thick exosporium, and red arrowheads indicate spores with a thin exosporium. Bottom panel shows the percentage of each exosporium thickness (blue bar: thick exosporium, red bar: thin exosporium). C) Appendage occurrence in unfractionated, upper fraction, and lower fraction spores observed by phase-contrast microscopy. The black arrowheads indicate spores with no appendages. Blue arrowheads correspond to type 1 appendage and red arrowheads to type 2 appendages. Percentage of appendage type is shown in the bottom panel (blue bar: type 1 appendages; red bar: type 2 appendages).

In the other hand, another characteristic that bifurcates the differentiation pathway in spore formation is the polar appendage, that can be present as a robust structure, a short appendage or be absent [14]. Additionally, short or absent appendage are related to poor germination in response to taurocholate [14]. Phase contrast microscopy shows polar appendage in two different morphologies: appendage type 1, a polar prolongation (Fig. 2C), and appendage type 2, diffuse and round prolongation. In unfractionated spores, only 6.4% of total spores exhibited appendage, being 3.6% type 1 and 2.8% type 2. Interestingly, appendage type 1 was enriched in the lower fraction (24.0% of spores) while appendage type 2 was enriched in the upper fraction (12.7% of spores). Collectively, these results provide evidence that thick exosporium and appendage type 1 can be enriched by a density gradient.

To assess whether these two spore-morphotypes could play a role in the adherence of spores to intestinal epithelial cells, we performed an infection assay of Caco-2 monolayers. Results show that adherence to epithelial cells is not different among fraction and is similar to unfractionated spores (Fig. S2). These results indicate that enrichment of appendages and thick-exosporium in the lower fraction does not affect spore-adherence to intestinal epithelial cells *in vitro*.

### Heterogenous distribution of spore coat and exosporium proteins detected by SNAP translational fusions in *C. difficile* R20291 spores

Prior results demonstrated that intrinsic variability is operating at the exosporium level, as evidenced by appendage variability and exosporium layer variability [10, 11]. To identify the specific distribution of spore coat and exosporium proteins, translational SNAP fusions were used. *C. difficile* R20291 cultures take 48 h in 70:30 agar plate to render 6 morphological stages of sporulation (Figure S3). 48-h sporulating culture incubated with SNAP reagent exhibited high heterogeneity in the pattern of distribution and intensity of fluorescence (Fig. S4). Statistical analysis of the fluorescence distribution revealed that none of these fusions followed a normal distribution (Shapiro-Wilk test, P < 0.0001) (Fig. S5), suggesting that two or more spore-populations might be present in each SNAP fusion proteins.

Patterns of distribution analysis of fluorescence shows that several fluorescence types can be found when coat and exosporium SNAP fusion proteins are examined (Fig. 3), when analyzing appendage and appendage-free spores in unfractionated spore population. Specifically, we observed that spores carrying CdeB-SNAP fusion exhibit only polar fluorescence signal, despite presence or absence of polar appendage (Fig. 3A). The cysteine rich protein CdeC-SNAP fusion shows homogeneous fluorescence around the spore when the appendage is absent in 54% of fluorescent spores, whereas positive-appendage spores had three types of fluorescence distribution at the poles (Fig. 3B). Notably, regardless of the absence of polar appendage in spores carrying CdeM-SNAP fusion, only half of fluorescent spores (52%) exhibited a homogeneous fluorescence distribution around the spore; while other fluorescence types were evidenced at one, both poles or at one pole-and-sides of the spore (Fig. 3C). Homogeneous fluorescence around spores was observed in 68% of appendage-negative spores carrying BclA3-SNAP fusion, while the rest exhibit polar-and-side fluorescence (Fig. 3D). Among spore coat protein-fusions, we observed that the spores carrying CotA-SNAP fusion had a regular fluorescence signal around the spore, and a small fraction had a strong fluorescence intensity in the spore pole where a type 2 appendage appears (Fig. 3E). By contrast, spores carrying CotB-, CotD- and CotE-SNAP fusions exhibited pole-located fluorescence signal (Fig. 3F, G, H).

**Figure 3.**
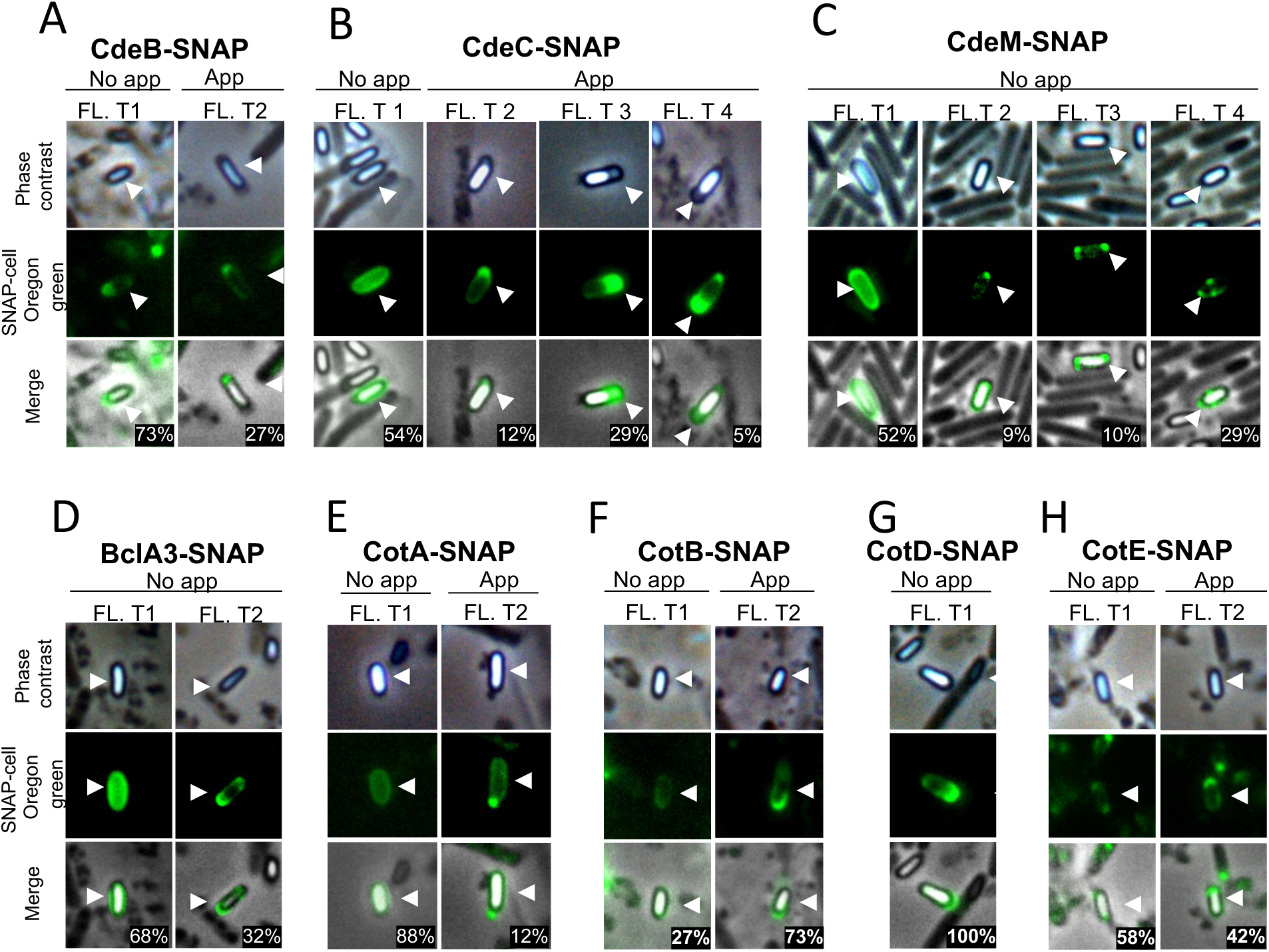
Fluorescence distribution of SNAP-spore-coat and -exosporium protein fusions in *C. difficile* spores. Localization of SNAP-fusions with exosporium proteins in *C. difficile* strain R20291. The strains expressing A) CdeB-, B) CdeC-, C) CdeM-, D) BclA3-, E) CotA-, F) CotB-, G) CotD- or H) CotE-SNAP fusion were grown for 48 hours in 70:30 agar plates and collected for labelling with Oregon green SNAP cell. Fl. t: fluorescence type. White arrows indicate the reference spore. The number in the panels are the percentage of spores found with specific. Numbers in merge image indicate the mean of the relative fluorescence intensity.

Since translational fusions of SNAP protein were cloned into plasmid pFT48, which yields 4-10 copy number per cell [6, 18], we assessed whether the overexpression of exosporium and spore coat proteins fused to the SNAP reporter protein would affect the formation of appendage (Fig. S6). Nearly 6% of the spores carrying the empty vector exhibited appendage. No significant increase in the percentage of appendage was evidenced in spores carrying CdeB (8% of total spores). However, spores carrying CdeC-, CotB- and CotE-SNAP fusion had an increased appendage-positive spores (Fig. S6). Expression of CdeM-, BclA3-, CotA-, and CotD-SNAP fusions led to completely lost of appendage (Fig. S6). Collectively, these results demonstrate that multicopy expression of SNAP fusion proteins differentially affects appendage formation in *C. difficile* spores.

### Effect of overexpression of the exosporium proteins CdeC, and CdeM on the ultrastucture of R20291 spore

Since exosporium protein CdeC and CdeM are key proteins for the proper assembly of the spore outer layers and appendage [8, 15], we evaluate if the extrachromosomal overexpression of translational SNAP fusions of these exosporium proteins (CdeC and CdeM) affect ultrastructural features of *C. difficile* spores. Both exosporium morphotypes (thin and thick) were observed in *C. difficile* R20291 carrying empty vector (Fig. 4A); thin exosporium is the most abundant morphotype (Fig 4A), while the thick morphotype is present in 30% of spores, consistent with previous reports [10]. Interestingly, overexpression of CdeC resulted in nearly 75% of the spores with a thick exosporium layer (Fig. 4A, B). CdeC-SNAP fusion also resulted in an amorphous accumulation of electron-dense material in the spore surface with bumps and hair-like projections (Fig. 4C); these spores correspond to nearly 28% of the analyzed spore sample (n = 100 spores). Expression of CdeM-SNAP fusion did not impact the proportion of exosporium morphotypes, however, *C. difficile* strain expressing CdeM-SNAP fusion produced spores with a disorganized and uncompact external material (Fig. 4C). This material seems to be loosely attached to the polar pole of the spore (Fig. 4C). Collectively, these results suggest that episomal expression of CdeC-SNAP fusion, but not CdeM-SNAP, affects the abundance and morphology of thick exosporium morphotype.

**Figure 4.**
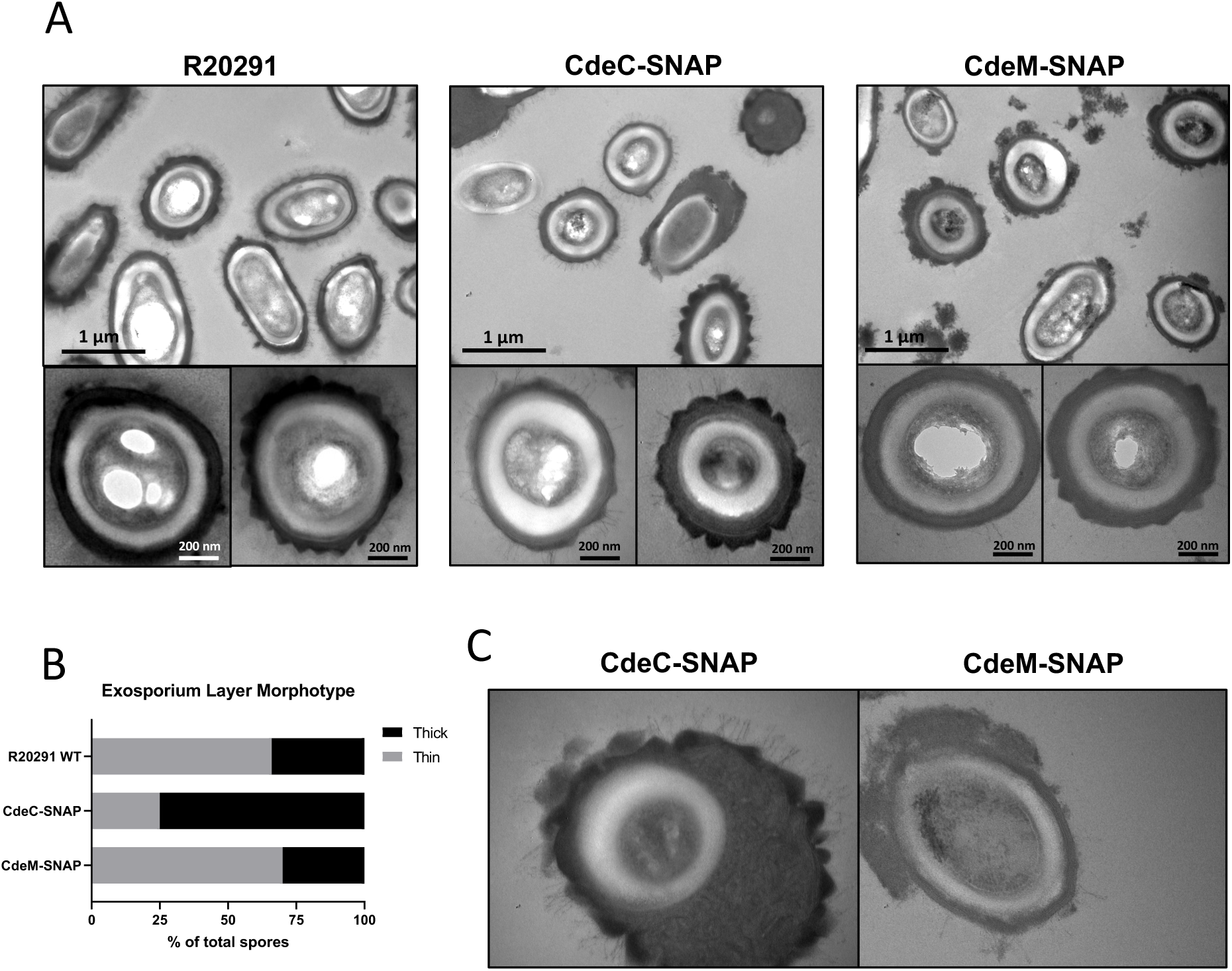
CdeC-SNAP fusions affects the exosporium ultrastructure of *C. difficile* R20291 spores. A) Purified spores of the CdeC-, CdeM-SNAP fusions were analyzed by TEM and the formation of exosporium morphotypes was evaluated. Aberrant structures are indicated by blue arrows. Bar scale: 1 μm and 200 nm. B) Exosporium morphotypes were quantified in each SNAP fusion. C) Spores carrying CdeC-SNAP spores exhibiting an accumulation of electron-dense material around the spore in one or two poles. Spores carrying CdeM-SNAP exhibit a disorganized exosporium and accumulation of material.

## Discussions

*C. difficile* spores are metabolically dormant life forms that are able to resist antibiotic exposure and act as transmission vectors, disseminating disease. The outermost layer of *C. difficile* spores is thought to act as the site of contact during the first stages of infection. Most epidemically relevant strains have an exosporium with hair-like projections that builds on top of a thin layer of electron dense material surrounding the spore coat; alternatively, these projections may also build on top of a thicker layer of electron dense material that surrounds the spore coats [10]. In this work, we provide more insight into the variability of these two types of exosporium layer of *C. difficile* spores.

A first finding of this work was that the different spore-morphotypes are formed simultaneously during sporulation. The simultaneous formation of both exosporium morphotypes (thin and thick) observed by transmission electron micrographs of sporulating cultures of R20291 in 70:30 as well as in TY agar plates suggests that the mechanism that drives each morphotype occurs independent of the culture conditions and in a subset of sporulating cells (Fig. 1). These observations also show that final morphotype is independent of sporulation timing (Fig. 1). Though it is intriguing that the small fraction of thick exosporium spores is maintained in different sporulating medium conditions. However, several questions arise; there is no genetic evidence of such a regulatory circuit that might regulate thickness of the outer layer of *C. difficile* spores. Potential candidates include the excision of the skin element in the mother cell by the recombinase gene *spoIVCA* which leads to active SigK and spores of *C. difficile* 630 strain with an electron dense exosporium layer [19]. The observation that thick exosporium layer spores are formed in aged sporulating cultures suggest that environmental conditions and/or quorum sensing factors, and/or additional unidentified stimuli might be implicated in the exosporium morphotype development. Further studies to identify and characterize the mechanisms that regulate the formation of thin and/or thick exosporium morphotypes are underway.

Several tools have been developed to track down proteins in anaerobic species such as *C. difficile* [17, 18]. Here we provide evidence that, there is great variability in the expression of SNAP reporter fusions with exosporium and spore coat proteins of *C. difficile*. We observed high variability in the fluorescence intensity and also in the distribution pattern of SNAP-fusion proteins. First, not all of the spores, exhibited fluorescence intensity, which could be attributed to poor folding of the SNAP fluorescent fusion protein or to low expression levels. In accordance to these observations, Imamura *et al*. (2011), also observed that not all of the spores expressing GFP-fusion proteins of the outer-most layer of *B. subtilis* spores yielded detectable fluorescence [20]. We also observed that the fluorescence distribution does not follow a normal distribution, and in most cases exhibited a skewed distribution, indicating that there might be subpopulation of spores with different spore surface protein compositions. Supporting this notion, we observed that the fluorescent distribution of CdeC-, CdeM-, BclA3- and CotA-SNAP fusions exhibited two distinctive patterns of fluorescence (Fig. 3): i) a homogenous distribution surrounding the spore; and ii) a defined pattern at one or both poles of the spore. Antunes *et al*. (2018) reported similar variability in the fluorescence pattern of SNAP-translational fusion with the exosporium protein, CdeC [14]. Results from Calderón-Romero *et al*. (2018) provide evidence that the absence of CdeC or CdeM affect the presence of BclA3, CdeB, CotA and CotB in strain 630 [8]. Notably, absence of CdeM affects the presence of CdeC but not vice versa in a 630 background [8], suggesting that the different fluorescence patterns could obey different assembly mechanisms. Studies to address these questions will provide insight into the mechanisms underlying the variability of the exosporium layer of *C. difficile* spores.

We were able to separate R20291 spores into two spore populations; one less dense fraction with similar abundance of thick exosporium layer spores and appendage-positive spores as unfractionated spores; and a denser fraction, enriched in thick exosporium layer spores and polar appendages. These observations are in agreement with the enrichment of appendage positive spores of strain 630 through density gradient [14]. Although, our results suggest that both phenotypes (i.e., thickness of the exosporium layer and the presence/absence of polar appendage) become enriched in the same gradient fraction, it is likely that the mechanism driving appendage formation and thick exosporium layer might be different. For example, absence of CdeM affects the thickness of the exosporium layer but not appendages presence [8, 14]. In our experimental conditions, we observed no difference in the adherence of spores from the upper and lower fractions to monolayers of epithelial Caco-2 cells. By contrast, Antunes *et al*. (2018), demonstrated that 630 spores enriched in appendage by density gradient germinated faster than fractions of spores with no appendage [14]. The exact role of these phenotypes in *C. difficile* pathogenesis is unclear and require further research.

Recent work has demonstrated that cysteine-rich proteins are responsible for the correct formation of the exosporium layer of *C. difficile* spores in several genetic backgrounds [8, 14, 21]. In R20291 and 630 strain, inactivation of *cdeC* leads to spores with an aberrantly assembled exosporium layer, and loss of various additional exosporium proteins, including CdeM. Similarly, in 630 strain, inactivation of *cdeM* also affects the assembly of the exosporium layer of *C. difficile* spores, but to a more superficial extent; that is, while *cdeC* inactivation affects the thickness of the spore coat and exosporium layer, *cdeM* inactivation only leads to loss of the exosporium layer. In this context, our results demonstrate that the multicopy expression of CdeC- and CdeM-SNAP fusion proteins lead to different effects on the surface of *C. difficile* spores (Fig. 4). Multicopy expression of CdeC-SNAP led to an increase in the ratio of thick-exosporium spores upon as well as an increase in spores with a amorphously thick exosporium layer (Fig. 4). It was also noteworthy to observe that the amorphous exosporium layer of these spores had electron-dense material deposited on the bumps formed and the presence of the typical hair-like projections observed in wild-type spores. This indicates that CdeC-SNAP fusion is functional, and that CdeC, is able to drive the formation of thick-exosporium layer. On the other hand, the CdeM-SNAP fusion, yielded spores with a similar ratio of thin/thick exosporium layer as spores carrying vector control, but with immunofluorescence signal through the entire spore as well as in one or both poles of the spores (Fig. 4). Multi-copy expression of CdeM-SNAP had no effect on the proportion of thin/thick exosporium ratio but led to an increased number of spores with a diffuse loose material partially surrounding the spore. Altogether these results contribute to understand the variability in the outermost layer of *C. difficile* spores and raise new questions about the underlying mechanisms of assembly and variability of this layer.

## 5. Acknowledgements

This work was supported by grants from Fondo Nacional de Ciencia y Tecnología de Chile (FONDECYT Grant 1151025, 1191601) and by Millennium Science Initiative of the Ministry of Economy, Development and Tourism to D.P-S. Support was also provided by a grant from Fondo de Fomento al Desarrollo Científico y Tecnológico (FONDEF) ID18|10230 to M.P-G and D.P-S.

## Supplementary Figures

**Figure S1. Transmission electron micrographs of 1-day- and 5-day-old sporulating cultures in TY agar plates**. A) Sporulating cultures in TY medium harvested after 1 day (left panel) and 5 days (right panel) of growth in TY medium. Cultures were fixed under anaerobic conditions prior to processing for transmission electron micrograph analysis. White arrow: thin exosporium spore, Black arrow: thick exosporium spore. Bar scale: 2 μm. B) Ultrastructural analysis of the exosporium layer. Thin and thick exosporium morphotype spores evidenced at day 1 (left panel) and day 5 (right panel). Scale bar: 200 nm. C) Percentage of thin and thick exosporium at day 1 (gray bar) and day 5 (black bar). Bar indicates standard error obtained with 3 different batch of culture fixed, processed and analyzed separately. N=50 spores. P-value indicated.

**Figure S2. Adherence of nycodenz-separated *C. difficile* R20291 spores to intestinal epithelial Caco-2 cell *in vitro***. A) Monolayers of undifferentiated 2-day-old Caco-2 and, B), differentiated 8-day-old Caco-2 cells were infected at a MOI of 10 with *C. difficile* R20291 unfractionated, upper and lower fractions spores for 3 h at 37°C. Unbound spores were removed by rinsing with PBS. Adherence was quantified as described in the Method section. Graphs represent the average of three independent experiments, error bars represent standard errors of the means. n.s.: no significance.

**Figure S3. Sporulation of *C. difficile* R20291**. A) Dynamics of sporulation. Different stages of the sporulation process: stage 1, vegetative cell; stage 2, asymmetric division with septum; stage 3, curvature in septum membrane; stage 4, conformed forespore; stage 5, pre-spore found as a bright structure; and stage 6, spore released. Membrane (FM4-64; red) and DNA (Hoechst; blue) stains are shown. White arrowhead indicates the cell or spore in the corresponding stage. Sporulation process is represented: mother cell membrane lined in blue, spore inner membrane in red and spore external layers in green. B) Quantification of total cell or spore of each stage in time (18, 24 and 48 h). Total cells for the quantification > 1800.

**Figure S4. SNAP fusions proteins in coat and exosporium of *C. difficile* R20291 spores**. Phase-contrast microscopy and fluorescence microscopy of *C. difficile* R20291 containing SNAP fusion, cultured by 48 hours in 70:30 agar plate and incubated with 250 nM of SNAP cell Oregon green (New England Biolabs®). White arrow indicates high fluorescence spores. Scale bar: 10 μm.

**Figure S5. Frequency distribution of SNAP proteins fusions of *C. difficile* R20291 spores**. *C-difficile* containing each SNAP fusion was independently incubated with 250 nM for 48 h with SNAP cell Oregon green (New England Biolabs®). A) CdeB-SNAP; B) CotA-SNAP; C) CdeC-SNAP; D) CotB-SNAP; E) CdeM-SNAP: F) CotD-SNAP; G) BclA3-SNAP; H) CotE-SNAP. n > 300. Shapiro-Wilk test was used and all sporulating cultures carrying SNAP fusions of exosporium and spore-coat proteins had P<0,0001.

**Figure S6. Presence or absence of appendage of *C. difficile* R20291 spores with SNAP fusions**. Percentage of *C. difficile* R20291 spores with empty vector and SNAP fusions are shown. The classification was made observing the spores in contrast phase microscopy of spores that carrying the SNAP fusions.

## References

1. Martin JS, Monaghan TM, Wilcox MH. *Clostridium difficile* infection: epidemiology, diagnosis and understanding transmission. Nat Rev Gastroenterol Hepatol. 2016;13(4):206-16. Epub 2016/03/10. doi: 10.1038/nrgastro.2016.25. PubMed PMID: 26956066.

2. Rupnik M, Wilcox MH, Gerding DN. Clostridium difficile infection: new developments in epidemiology and pathogenesis. Nat Rev Microbiol. 2009;7(7):526-36. Epub 2009/06/17. doi: 10.1038/nrmicro2164. PubMed PMID: 19528959.

3. Lessa FC, Mu Y, Bamberg WM, Beldavs ZG, Dumyati GK, Dunn JR, et al. Burden of *Clostridium difficile* infection in the United States. N Engl J Med. 2015;372(9):825-34. Epub 2015/02/26. doi: 10.1056/NEJMoa1408913. PubMed PMID: 25714160.

4. Deakin LJ, Clare S, Fagan RP, Dawson LF, Pickard DJ, West MR, et al. The *Clostridium difficile spo0A* gene is a persistence and transmission factor. Infection and Immunity. 2012;80(8):2704–11. doi: 10.1128/iai.00147-12.

5. Fimlaid KA, Bond JP, Schutz KC, Putnam EE, Leung JM, Lawley TD, et al. Global analysis of the sporulation pathway of *Clostridium difficile*. PLoS Genet. 2013;9(8):e1003660. Epub 2013/08/21. doi: 10.1371/journal.pgen.1003660. PubMed PMID: 23950727; PubMed Central PMCID: PMCPMC3738446.

6. Pereira FC, Saujet L, Tome AR, Serrano M, Monot M, Couture-Tosi E, et al. The spore differentiation pathway in the enteric pathogen *Clostridium difficile*. PLoS Genet. 2013;9(10):e1003782. Epub 2013/10/08. doi: 10.1371/journal.pgen.1003782. PubMed PMID: 24098139; PubMed Central PMCID: PMCPMC3789829.

7. Mora-Uribe P, Miranda-Cardenas C, Castro-Cordova P, Gil F, Calderon I, Fuentes JA, et al. Characterization of the Adherence of *Clostridium difficile* Spores: The Integrity of the Outermost Layer Affects Adherence Properties of Spores of the Epidemic Strain R20291 to Components of the Intestinal Mucosa. Front Cell Infect Microbiol. 2016;6:99. Epub 2016/10/08. doi: 10.3389/fcimb.2016.00099. PubMed PMID: 27713865; PubMed Central PMCID: PMCPMC5031699.

8. Calderón-Romero P, Castro-Córdova P, Reyes-Ramírez R, Milano-Céspedes M, Guerrero-Araya E, Pizarro-Guajardo M, et al. Clostridium difficile exosporium cysteine-rich proteins are essential for the morphogenesis of the exosporium layer, spore resistance, and affect *C. difficile* pathogenesis. PLOS Pathogens. 2018;14(8):e1007199. doi: 10.1371/journal.ppat.1007199.

9. Paredes-Sabja D, Sarker MR. Adherence of *Clostridium difficile* spores to Caco-2 cells in culture. Journal of Medical Microbiology. 2012;61(9):1208–18. doi: doi:10.1099/jmm.0.043687-0.

10. Pizarro-Guajardo M, Calderon-Romero P, Paredes-Sabja D. Ultrastructure Variability of the Exosporium Layer of *Clostridium difficile* Spores from Sporulating Cultures and Biofilms. Appl Environ Microbiol. 2016;82(19):5892-8. Epub 2016/07/31. doi: 10.1128/AEM.01463-16. PubMed PMID: 27474709; PubMed Central PMCID: PMCPMC5038037.

11. Pizarro-Guajardo M, Calderon-Romero P, Castro-Cordova P, Mora-Uribe P, Paredes-Sabja D. Ultrastructural Variability of the Exosporium Layer of *Clostridium difficile* Spores. Appl Environ Microbiol. 2016;82(7):2202-9. Epub 2016/02/07. doi: 10.1128/AEM.03410-15. PubMed PMID: 26850296; PubMed Central PMCID: PMCPMC4807528.

12. Díaz-González F, Milano M, Olguin-Araneda V, Pizarro-Cerda J, Castro-Córdova P, Tzeng S-C, et al. Protein composition of the outermost exosporium-like layer of *Clostridium difficile* 630 spores. Journal of Proteomics. 2015;123:1–13.

13. Pizarro-Guajardo M, Olguin-Araneda V, Barra-Carrasco J, Brito-Silva C, Sarker MR, Paredes-Sabja D. Characterization of the collagen-like exosporium protein, BclA1, of *Clostridium difficile* spores. Anaerobe. 2014;25:18-30. Epub 2013/11/26. doi: 10.1016/j.anaerobe.2013.11.003. PubMed PMID: 24269655.

14. Antunes W, Pereira FC, Feliciano C, Saujet L, Vultos Td, Couture-Tosi E, et al. Structure and assembly of a *Clostridioides difficile* spore polar appendage. bioRxiv. 2018:468637. doi: 10.1101/468637.

15. Barra-Carrasco J, Olguin-Araneda V, Plaza-Garrido A, Miranda-Cardenas C, Cofre-Araneda G, Pizarro-Guajardo M, et al. The *Clostridium difficile* exosporium cysteine (CdeC)-rich protein is required for exosporium morphogenesis and coat assembly. J Bacteriol. 2013;195(17):3863-75. Epub 2013/06/25. doi: 10.1128/JB.00369-13. PubMed PMID: 23794627; PubMed Central PMCID: PMCPMC3754587.

16. Paredes-Sabja D, Cofre-Araneda G, Brito-Silva C, Pizarro-Guajardo M, Sarker MR. Clostridium difficile spore-macrophage interactions: spore survival. PLoS One. 2012;7(8):e43635. Epub 2012/09/07. doi: 10.1371/journal.pone.0043635. PubMed PMID: 22952726; PubMed Central PMCID: PMCPMC3428350.

17. Cassona CP, Pereira F, Serrano M, Henriques AO. A Fluorescent Reporter for Single Cell Analysis of Gene Expression in *Clostridium difficile*. Methods Mol Biol. 2016;1476:69-90. Epub 2016/08/11. doi: 10.1007/978-1-4939-6361-4_6. PubMed PMID: 27507334.

18. Ransom EM, Ellermeier CD, Weiss DS. Use of mCherry Red Fluorescent Protein for Studies of Protein Localization and Gene Expression in *Clostridium difficile*. Applied and Environmental Microbiology. 2015;81(5):1652–60. doi: 10.1128/aem.03446-14.

19. Serrano M, Kint N, Pereira FC, Saujet L, Boudry P, Dupuy B, et al. A Recombination Directionality Factor Controls the Cell Type-Specific Activation of sigmaK and the Fidelity of Spore Development in *Clostridium difficile*. PLoS Genet. 2016;12(9):e1006312. Epub 2016/09/16. doi: 10.1371/journal.pgen.1006312. PubMed PMID: 27631621; PubMed Central PMCID: PMCPMC5025042.

20. Imamura D, Kuwana R, Takamatsu H, Watabe K. Proteins involved in formation of the outermost layer of *Bacillus subtilis* spores. J Bacteriol. 2011;193(16):4075-80. Epub 2011/06/15. doi: 10.1128/JB.05310-11. PubMed PMID: 21665972; PubMed Central PMCID: PMCPMC3147665.

21. Barra-Carrasco J, Olguín-Araneda V, Plaza-Garrido Á, Miranda-Cárdenas C, Cofré-Araneda G, Pizarro-Guajardo M, et al. The *Clostridium difficile* Exosporium Cysteine (CdeC)-Rich Protein Is Required for Exosporium Morphogenesis and Coat Assembly. Journal of Bacteriology. 2013;195(17):3863-75. Epub 2013/06/25. doi: 10.1128/JB.00369-13. PubMed PMID: PMC3754587; PubMed Central PMCID: PMCPMC3754587.

