## Supplementary material for "Characterization of exosporium layer variability of *Clostridioides difficile* spores in the epidemically relevant strain R20291": Supp Fig

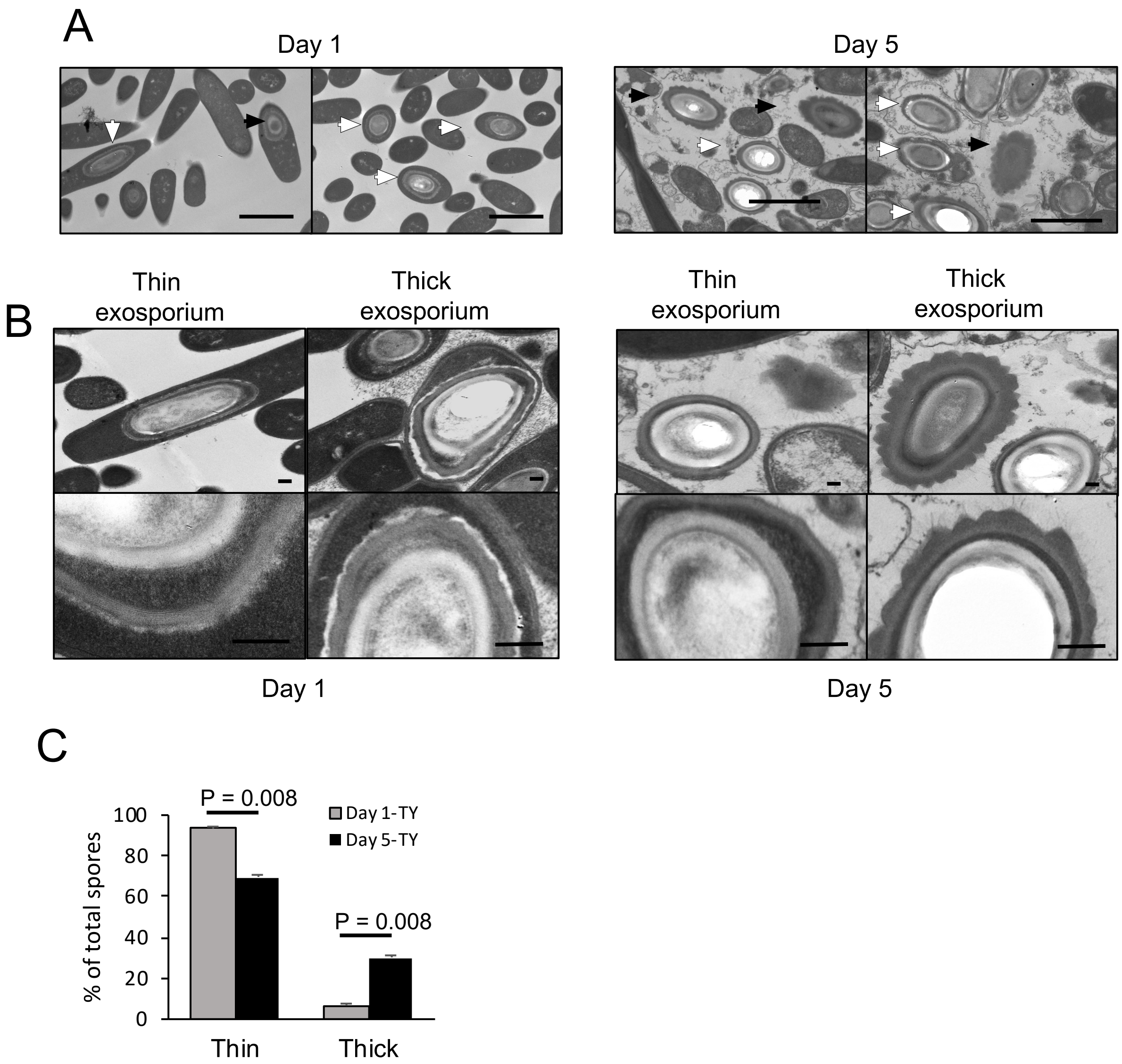


**Figure S1. Transmission electron micrographs of 1-day- and 5-day-old sporulating cultures in TY agar plates.** A) Sporulating cultures in TY medium harvested after 1 day (left panel) and 5 days (right panel) of growth in TY medium. Cultures were fixed under anaerobic conditions prior to processing for transmission electron micrograph analysis. White arrow: thin exosporium spore, Black arrow: thick exosporium spore. Bar scale: 2 µm. B) Ultrastructural analysis of the exosporium layer. Thin and thick exosporium morphotype spores evidenced at day 1 (left panel) and day 5 (right panel). Scale bar: 200 nm. C) Percentage of thin and thick exosporium at day 1 (gray bar) and day 5 (black bar). Bar indicates standard error obtained with 3 different batch of culture fixed, processed and analyzed separately. N=50 spores. P-value indicated.

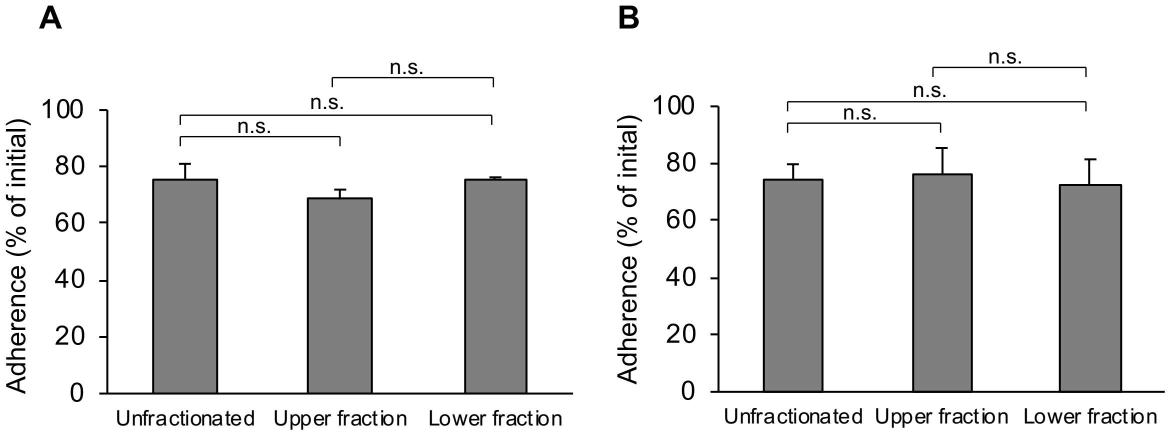


**Figure S2. Adherence of nycodenz-separated *C. difficile* R20291 spores to intestinal epithelial Caco-2 cell *in vitro*.** A) Monolayers of undifferentiated 2-day-old Caco-2 and, B), differentiated 8-day-old Caco-2 cells were infected at a MOI of 10 with *C. difficile* R20291 unfractionated, upper and lower fractions spores for 3 h at 37ºC. Unbound spores were removed by rinsing with PBS. Adherence was quantified as described in the Method section. Graphs represent the average of three independent experiments, error bars represent standard errors of the means. n.s.: no significance.

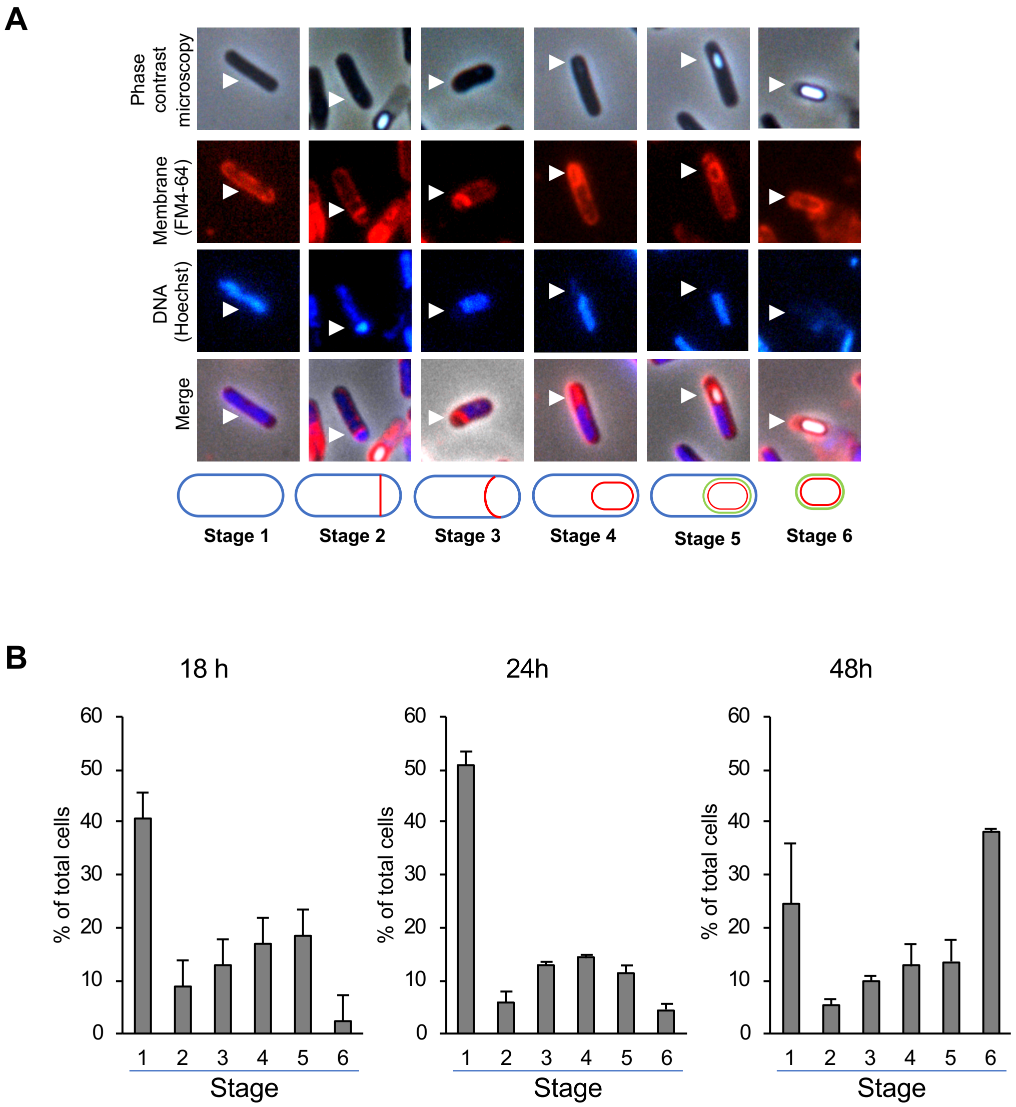


**Figure S3. Sporulation of *C. difficile* R20291.** A) Dynamics of sporulation. Different stages of the sporulation process: stage 1, vegetative cell; stage 2, asymmetric division with septum; stage 3, curvature in septum membrane; stage 4, conformed forespore; stage 5, pre-spore found as a bright structure; and stage 6, spore released. Membrane (FM4-64; red) and DNA (Hoechst; blue) stains are shown. White arrowhead indicates the cell or spore in the corresponding stage. Sporulation process is represented: mother cell membrane lined in blue, spore inner membrane in red and spore external layers in green. B) Quantification of total cell or spore of each stage in time (18, 24 and 48 h). Total cells for the quantification > 1800.

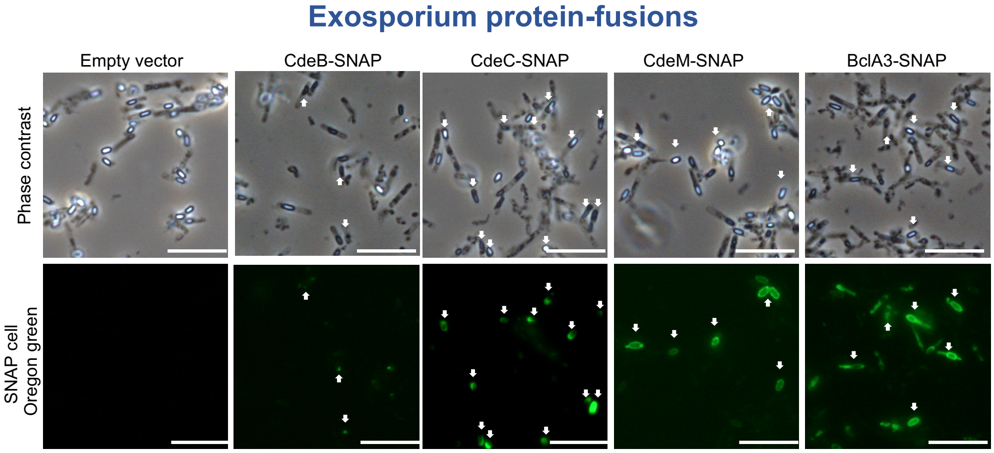


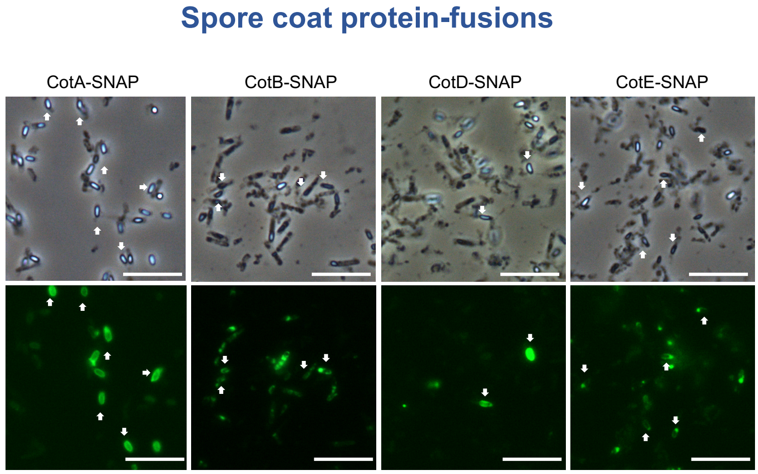


**Figure S4. SNAP fusions proteins in coat and exosporium of *C. difficile* R20291 spores.** Phase-contrast microscopy and fluorescence microscopy of *C. difficile* R20291 containing SNAP fusion, cultured by 48 hours in 70:30 agar plate and incubated with 250 nM of SNAP cell Oregon green (New England Biolabs®). White arrow indicates high fluorescence spores. Scale bar: 10 μm.

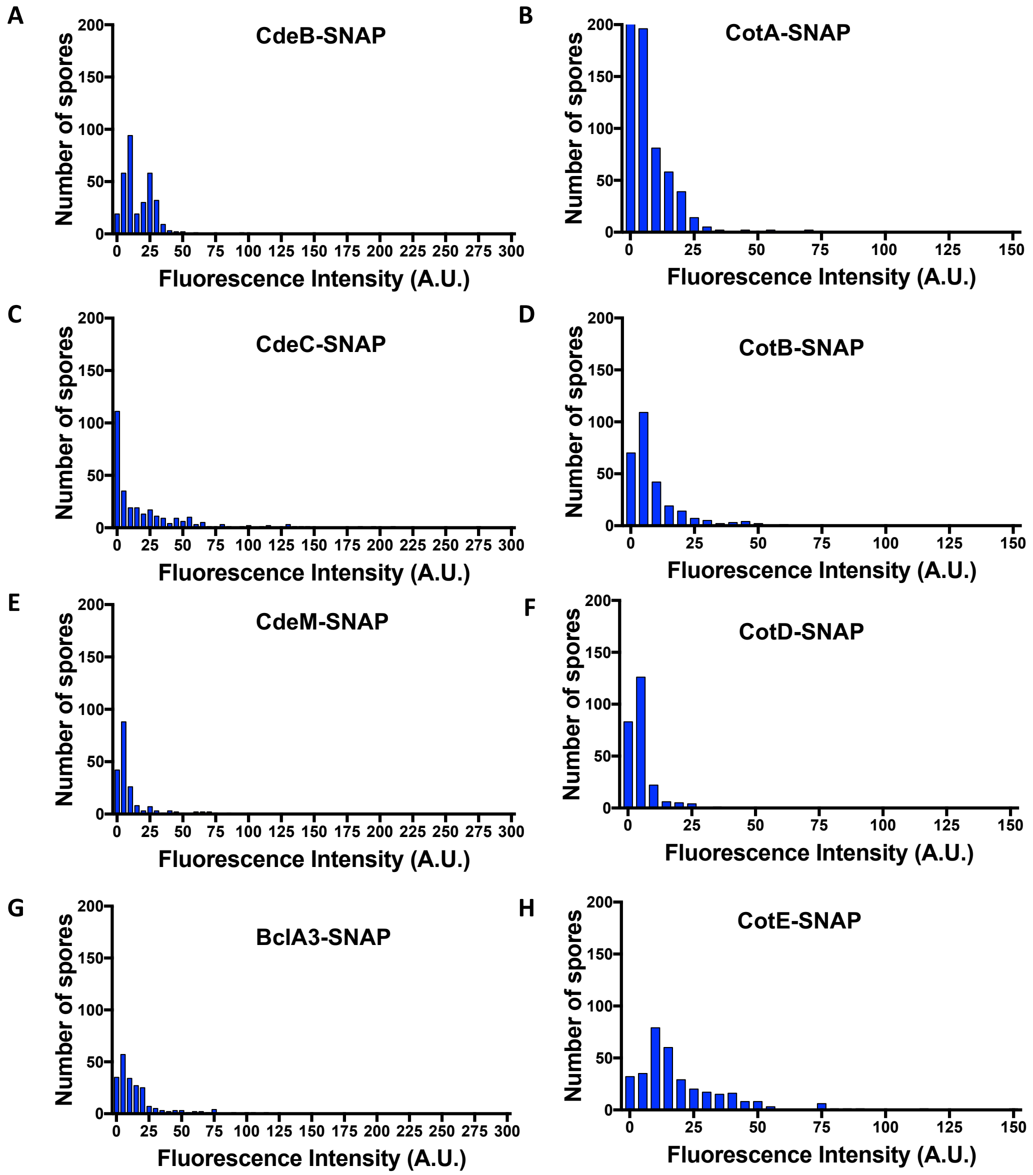


**Figure S5. Frequency distribution of SNAP proteins fusions of *C. difficile* R20291 spores.** Culture with each SNAP fusions was independently incubated with 250 nM for 48 h with SNAP cell Oregon green (New England Biolabs®). A) CdeB-SNAP; B) CotA-SNAP; C) CdeC-SNAP; D) CotB-SNAP; E) CdeM-SNAP: F) CotD-SNAP; G) BclA3-SNAP; H) CotE-SNAP. n > 300. Shapiro-Wilk test was used and all sporulating cultures carrying SNAP fusions of exosporium and spore-coat proteins had P<0,0001.

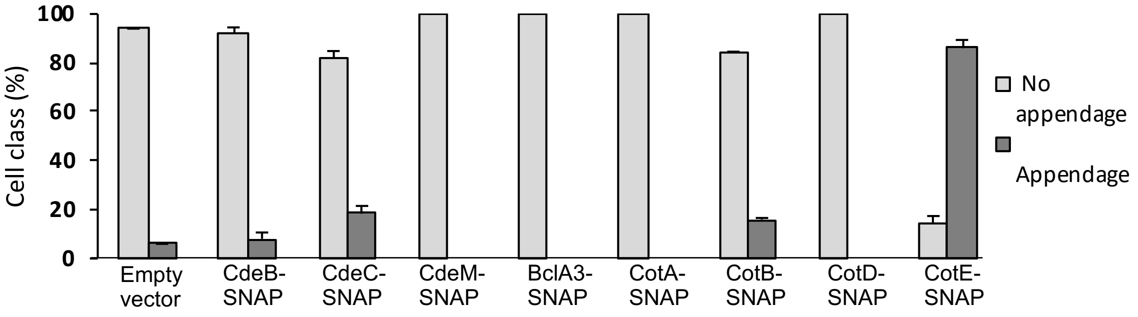


**Figure S6. Presence or absence of appendage of *C. difficile* R20291 spores with SNAP fusions.** Percentage of *C. difficile* R20291 spores with empty vector and SNAP fusions are shown. The classification was made observing the spores in contrast phase microscopy of spores that carrying the SNAP fusions.
