## Supplementary material for "Characterization of exosporium layer variability of *Clostridioides difficile* spores in the epidemically relevant strain R20291": Supp table

**Table S1. Primers used in this work to create SNAP fusions.**

| Gene Locus[a] | Primer name | Primer sequence[b] | Position[c] |
|---|---|---|---|
| *cdeC* (CDR20291_0926) | P499 FP-cdeC-0926-EcoRI | GACGAATTCAATGAAATACGGGAGACCGTGTCTGG | -318 to -293 |
| | P500 RP-cdeC-0926-BamHI | GACGGATCCTCTGTGGCAACTTGGCTTTCC | +1195 to +1218 |
| *cdeM* (CDR20291_1478) | P562 FP-cdeM-1478-EcoRI | GACGAATTCCTGAATATGAAATAAACAACAGTTTATGTC | - 199 to -169 |
| | P563 RP-cdeM-1478-BamHI | GACGGATCCTTTTCTACAGCAGTTACAATTACATTTATG | +466 to 492 |
| *bclA3* (CDR20291_3193) | P508 FP-bclA3-3193-EcoRI | GACGAATTCGCTAAAGAGTACGGGCTGATTG | -1524 to -1503 |
| | P509 RP-bclA3-3193-BamHI | GACGGATCCATTTATTGCAATTCCTGCACTTG | +2011 to +2034 |
| *cotA* (CDR20291_1511) | P511 FP-cotA-1511-EcoRI | GACGAATTCTACTTCCTGATGTTGGTGTTAATATGCCA | -484 to -456 |
| | P512 RP-cotA-1511-BamHI | GACGGATCCTGCAATATAATCTATAGAATCTACACATACAAAG | + 891 to +924 |
| *cdeB* (CDR20291_2642) | P597 FP-cdeB-2642-EcoRI | TATATGAATTCCATACTCAAATTCTTCATCATC | -422 to -401 |
| | P598 RP-cdeB-2642-BamHI | GACGGATCCGTTAAGATTTCCTGCTTTATTAG | +641 to +663 |
| *cotB* (CDR20291_1360) | P599 FP-cotB-1360-EcoRI | TAATGAATTCCTTGAAATTGTTTGCATACTTAATT | -563 to -539 |
| | P600 RP-cotB-1360-BamHI | GACGGATCCCATGTTTTTATAACTCTCCAATATTC | +886 to +912 |
| *cotE* (CDR20291_1282) | P601 FP-cotE-1282-EcoRI | TATCAGAATTCCATGGAGATAACTAAAATTCTTAG | -268 to -245 |
| | P602 RP-cotE-1282-BamHI | GACGGATCCGAATTGCCCATAAATACCTTCAAGTTC | +2101 to +2128 |
| *cotD* (CDR20291_0523) | P603 FP-cotD-0523-EcoRI | CAGAAAGAATTCGCACAGAAAAAAGAGTAGG | -978 to -950 |
| | P604 RP-cotD-0523-BamH1 | TACGGATCCGAACTGATGCTTACACTC | +550 to +570 |

[a] Data base: EnsemblGenomes Gene, NCBI FN545816_R20291.
b Restriction site is underlined.
[c] The nucleotide position number begins from the first codon and refers to the relevant position within the respective gene sequence.

**Table S2. Plasmids and strains used in this work**

| Plasmid or strain name | Description | Reference |
|---|---|---|
| pFT58 | Derived from pMTL84121, contains SNAP sequence lacking start codon, between BamHI/HindIII sites. | (Pereira et al. 2013) |
| pPCR15 | CdeC-SNAP translational fusion. A 1553 bp PCR fragment amplified with P499/P500, cloned in BamHI/EcoRI sites of pFT58. | This work |
| pPCR16 | CdeM-SNAP translational fusion. A 1131 bp PCR fragment amplified with P562/P563, cloned in BamHI/EcoRI sites of pFT58. | This work |
| pPCR10 | BclA3-SNAP translational fusion. A 3683 bp PCR fragment amplified with P508/P509, cloned in BamHI/EcoRI sites of pFT58. | This work |
| pPCR11 | CotA-SNAP translational fusion. A 1107 bp PCR fragment amplified with P511/P512, cloned in BamHI/EcoRI sites of pFT58. | This work |
| pARR3 | CdeB-SNAP translational fusion. A 1090 bp PCR fragment amplified with P597/P598, cloned in BamHI/EcoRI sites of pFT58. | This work |
| pARR2 | CotB-SNAP translational fusion. A 1480 bp PCR fragment amplified with P599/P600, cloned in BamHI/EcoRI sites of pFT58. | This work |
| pARR6 | CotE-SNAP translational fusion. A 2400 bp PCR fragment amplified with P601/P602, cloned in BamHI/EcoRI sites of pFT58. | This work |
| pARR5 | CotD-SNAP translational fusion. A 1554 pb PCR fragment amplified with P603/P604, cloned in BamHI/EcoRI sites of pFT58. | This work |
| **Strain** | **Characteristics** | **Reference** |
| *C. difficile* R20201 | Ribotype 027, epidemically relevant strain | (McEllistrem et al. 2005) |
| *C. difficile* R20291 (pPCR15) | *C. difficile* R20291 carrying *cdeC*-SNAP fusions | This work |
| *C. difficile* R20291 (pPCR16) | *C. difficile* R20291 carrying *cdeM*-SNAP fusions | This work |
| *C. difficile* R20291 (pPCR11) | *C. difficile* R20291 carrying *cotA*-SNAP fusions | This work |
| *C. difficile* R20291 (pPCR10) | *C. difficile* R20291 carrying *bclA3*-SNAP fusions | This work |
| *C. difficile* R20291 (pARR6) | *C. difficile* R20291 carrying *cotE*-SNAP fusions | This work |
| *C. difficile* R20291 (pARR5) | *C. difficile* R20291 carrying *cotD*-SNAP fusions | This work |
| *C. difficile* R20291 (pARR2) | *C. difficile* R20291 carrying *cotB*-SNAP fusions | This work |
| *C. difficile* R20291 (pARR3) | *C. difficile* R20291 carrying *cdeB*-SNAP fusions | This work |
| *C. difficile* R20291 (pFT58) | *C. difficile* R20291 carrying empty vector | (Pereira et al. 2013) |
